# Client processing is altered by novel myopathy-causing mutations in the HSP40 J domain

**DOI:** 10.1101/2020.02.14.949792

**Authors:** Melanie Y. Pullen, Conrad C. Weihl, Heather L. True

## Abstract

The misfolding and aggregation of proteins is often implicated in the development and progression of degenerative diseases. Heat shock proteins (HSPs), such as the ubiquitously expressed Type II Hsp40 molecular chaperone, DNAJB6, assist in protein folding and disaggregation. Historically, mutations within the DNAJB6 G/F domain have been associated with Limb-Girdle Muscular Dystrophy type 1D, now referred to as LGMDD1, a dominantly inherited degenerative disease. Recently, novel mutations within the J domain of DNAJB6 have been reported in patients with LGMDD1. Since novel myopathy-causing mutations in the Hsp40 J domain have yet to be characterized and both the function of DNAJB6 in skeletal muscle and the clients of this chaperone are unknown, we set out to assess the effect of these mutations on chaperone function using the genetically tractable yeast system. The essential yeast Type II Hsp40, Sis1, is homologous to DNAJB6 and is involved in the propagation of yeast prions. Using phenotypic, biochemical, and functional assays we found that homologous mutations in the Sis1 J domain differentially alter the processing of specific yeast prion strains, as well as a non-prion substrate. These data suggest that the newly-identified mutations in the J domain of DNAJB6 cause aberrant chaperone function that leads to the pathogenesis in LGMDD1.

## Introduction

Molecular chaperones preserve protein homeostasis (1). A deficient chaperone network may lead to protein misfolding and aggregation often associated with protein conformational disorders such as Alzheimer’s Disease, Charcot-Marie-Tooth disease, distal hereditary motor neuropathies, and Limb Girdle Muscular Dystrophy, among others (2,3). Limb Girdle Muscular Dystrophy type 1D (LGMD1D), more recently termed LGMDD1 (4), is a disease characterized by proximal muscle weakness with moderate progression mediated by defective chaperone function (5). Historically, dominantly inherited disease-associated mutations in the type II Hsp40 co-chaperone DNAJB6 have been found within a 12 amino acid region known as the G/F domain (6–11). Recently, three novel pathogenic mutations associated with LGMDD1 have been identified within the J domain of DNAJB6 (12,13).

Since molecular chaperones are highly conserved from yeast to mammals, we used a yeast model system to study these disease-associated mutations (14–16). The essential yeast Type II Hsp40, Sis1, is homologous to DNAJB6 and plays an important role in yeast for the propagation of two prions, [*RNQ*^+^] and [*PSI*^+^] (17–22). Prions in yeast are epigenetic elements that form when the prion proteins form amyloid aggregates. These prion protein aggregates are propagated by fragmentation into propagons, which are then passed on to progeny during mitosis (14). Somewhat counterintuitively, chaperones not only promote prion propagation in yeast, but are essential for it (14,23–26). At the heart of this chaperone-mediated process is the Hsp40 Sis1, with the interaction between Hsp40 and Hsp70 chaperones being crucial for prion propagation in yeast (24,27–32). When chaperone function is defective, prion propagation is impaired (17,33–36). Thus, the loss of prion propagation provides a read-out for chaperone dysfunction in yeast.

Homologous LGMDD1-associated mutations of the DNAJB6 G/F domain impair the propagation of various yeast prion strains when expressed in Sis1 (37). We were inspired by this model to assess the effect of homologous J domain mutants in Sis1 on chaperone function. Fortunately, the yeast model system allows for a wide array of phenotypic, biochemical, and functional assays to assess the effect of these mutations on chaperone function (14,38). As such, we assessed the ability of the disease-associated mutant chaperone to process two specific clients of Sis1, Sup35 and Rnq1, which form the prion elements [*PSI*^+^] and [*RNQ*^+^], respectively. In addition, we performed functional assays with a non-prion client of the yeast chaperone machinery, firefly luciferase (FFL) (39). In conjunction, these assays elucidated nuances in client processing due to disease-associated mutations in a manner that would not be possible by assessing a single client protein or using a limited number of assays.

Here, we present evidence that disease-associated mutations in the J domain cause functional defects that differentially impair processing of native proteins in a client and conformer specific manner. Our work suggests that mutations in the J domain are responsible for altered protein processing and, potentially, disease pathogenesis.

## Materials and methods

### Protein sequence alignment structure modeling

Protein sequences for DNAJB6b orthologs were found on UniProt and subsequently aligned to homologous disease-causing J domain mutations in Sis1. Sequences were visualized using BioEdit (77) with multiple sequences aligned through the use of ClustalW2 (77–79). Mutant DNAJB6b structures were generated through the use of using I-TASSER (80–82) and visualized using PyMOL (83). Structural comparisons and protein threading of variants to wild-type protein structures was performed using wild-type Sis1 (PBD: 4RWU) (21).

### Yeast cultures, transformations and treatments

All experiments were performed in derivates of 74-D694 (*ade1-14 his3-Δ200 leu2-3,112 trp1-289 ura3-52)*. Yeast strains are [*PSI^+^*], [*psi*−], [*RNQ^+^*], or [*rnq*−], kindly gifted by S. Liebman (50,84) and J. Weissman (51). Yeast were cultured using YPD (1% yeast extract, 2% peptone, 2% dextrose), 1/4 YPD (0.25% yeast extract, 2% peptone, 2% dextrose) or synthetic defined (SD) media (0.67% yeast nitrogen base without amino acids, 2% dextrose) lacking specific amino acids using standard techniques. In order to study the function of *sis1* mutants in the absence of wild type *SIS1*, previously described plasmid shuffle strains carrying pRS316-SIS1 (37) were transformed with pRS314 (85) carrying *sis1* mutants. We selected for colonies that lost wild type *SIS1* through plasmid shuffle on plates containing 5-fluoroorotic acid (5-FOA). Plasmid transformations were performed by the PEG/LiOAC method (86). Plasmid pRS316-Sis1 was a kind gift from E. Craig (17,18). Plasmid 316-GPD-Lux was a kind gift from J. Weissman (67). Other plasmids are described below and were constructed using standard molecular techniques. For curing of prion strains, yeast were passaged twice on 3mM guanidine hydrochloride (GdnHCl) plates, then grown on complete media without GdnHCl for use in assays.

### Plasmid construction

Oligonucleotides used for site-directed mutagenesis are listed in S1 Table. Using pRS314-SIS1, the J domain LGMDD1 mutations were created by site-directed mutagenesis using the Agilent QuikChange II XL Site-Directed Mutagenesis Kit, as per manufacturer’s instructions along with the following oligonucleotides: 1890 and 1891 (S49V), 1892 and 1893 (E53A), 1894 and 1895 (N56L). Primer sequences were generated using Agilent’s online primer design program. Mutagenesis was confirmed by sequencing the entire coding region of *SIS1*.

### Yeast phenotypic assays

For yeast spottings, cultures were grown overnight in selective media. Cultures were pelleted, washed, and suspended in water to an optical density of 0.1. The normalized yeast solutions were pipetted into a 96-well plate, and serial dilutions (1:5) were made using a multichannel pipette. Yeast were spotted onto plates using an ethanol-sterilized 48-pin replicator. To monitor cell growth or the [*PSI*^+^] status of yeast cells with the spread plate assay, overnight cultures were normalized by A600, serially diluted 5-fold, and spotted on the indicated media. [*PSI*^+^] status was assessed by colony color on large 145mm plates in rich 1⁄4 YPD media. Plates were incubated for 5 days at 30°C followed by overnight incubation at 4°C for additional color development.

### Protein analysis

For boiled gel assays, yeast strains were cultured overnight. Cells were lysed with glass beads in buffer (25mM Tris-HCl pH7.5, 100mM NaCl, 1mM EDTA, protease inhibitors) and pre-cleared at 6,000 rpm for 1 minute at 4°C. Protein concentration of cell lysates was normalized using a Bradford assay and mixed with SDS-Page sample buffer (200mM Tris-HCl pH 6.8, 4% SDS, 0.4% bromophenol blue, 40% glycerol). Samples remained un-boiled and were loaded on a 12% polyacrylamide gel and run under constant current of 110V until the dye front migrated halfway through the resolving gel. The current was then stopped and, with the assembled gel remaining intact within the glass plates, they were sealed in plastic and boiled upright for 15mins in a 95-100°C water bath. After boiling, gels were removed from the plastic cover and were reinserted in the SDS-PAGE apparatus, where voltage was re-applied until the dye migrated to the bottom of the gel. SDS-PAGE was followed by standard western blotting with Sup35, Sis1, and Pgk1 antibodies. Sedimentation analysis for Rnq1 was performed as previously described (37,48). Semi-denaturing agarose gel electrophoresis (SDD-AGE) for [*PSI*^+^] was performed as previously described (37,87).

### Luciferase refolding

Luciferase refolding assays were performed as previously described (68). To monitor the ability of Sis1 to enhance refolding of luciferase *in vivo*, [*PSI*^+^] or [*rnq*−] yeast strains were transformed with the plasmid pRS316-GPD-Lux. Cultures were grown overnight in selective media and back diluted to an optical density of 0.3 in 8mL plastic culture tubes. Subsequently, cyclohexamide was added to a final concentration of 10ug/mL. Treated samples were then subjected to heat shock at 42°C for 22 minutes. Meanwhile, control samples were plated in 98-well clear bottom plates and kept at 30°C. After heat shock, 200ul of each sample was plated in triplicate on 98-well clear bottom plates. All cultures were allowed to recover at 30°C. Luminometer readings were taken at 30, 60, 90 and, 120 minutes. For data analysis, each triplicate was averaged. 2way ANOVA was performed using GraphPad Prism version 8.1.1 for Windows, GraphPad Software, La Jolla California USA, www.graphpad.com.

### Statistical methods

Data are reported as the means SEM of at least three independent experiments. Comparisons between multiple groups were performed by one-way analysis of variance test (ANOVA). For comparison between two groups, the paired t-test was used. In all cases, p<0.05 was considered significant. GraphPad Prism 8.1.1 software (GraphPad Software) was used for analysis.

### Antibodies and reagents

The following antibodies were used for Western Blots: rabbit anti-Sup35, dilution 1:1000 (True lab); rabbit anti-Sis1, dilution 1:5000 (Cosmo Bio, COP-080051); rabbit anti-Rnq1-11, dilution 1:2500 (True lab); mouse anti-Pgk1, dilution 1:2000 (Abcam, ab113687). For the firefly luciferase refolding assay, D-Luciferin (Sigma, L9504) was used as a substrate.

## Results

### Homologous mutations in the Sis1 J domain are conserved and potentially alter J domain and G/F domain interaction

Recently, novel mutations located within a 7 amino acid region of the DNAJB6 J domain have been reported to cause dominant distal myopathy (12,13). These variants were identified in five different families presenting with both distal and proximal muscle weakness. Initial studies examined these variants in patient tissue and cell culture studies in an attempt to elucidate whether these mutations lead to functional defects in DNAJB6’s anti-aggregation activity. From these studies, the recently identified DNAJB6 variants DNAJB6-A50V, DNAJB6-E54A and DNAJB6-S57L have been categorized as pathogenic, likely pathogenic, and variant of unknown significance, respectively (12).

To further examine the effect of these disease-associated variants on myopathy, we performed protein threading to identify homologous mutations in the yeast Hsp40 Sis1. Our comparative modeling identified homologous J domain mutations in Sis1 to be Sis1-S49V (homologous to DNAJB6-A50V), Sis1-E53A (homologous to DNAJB6-E54A), and Sis1-N56L (homologous to DNAJB6-S57L) (Fig 1A). In addition, our structural model recapitulates the observation that mutations in the J domain have the potential to disrupt important intramolecular interactions previously shown to stabilize the interface between the G/F and J domains (Fig 1B) (12,21).

**Fig 1.**
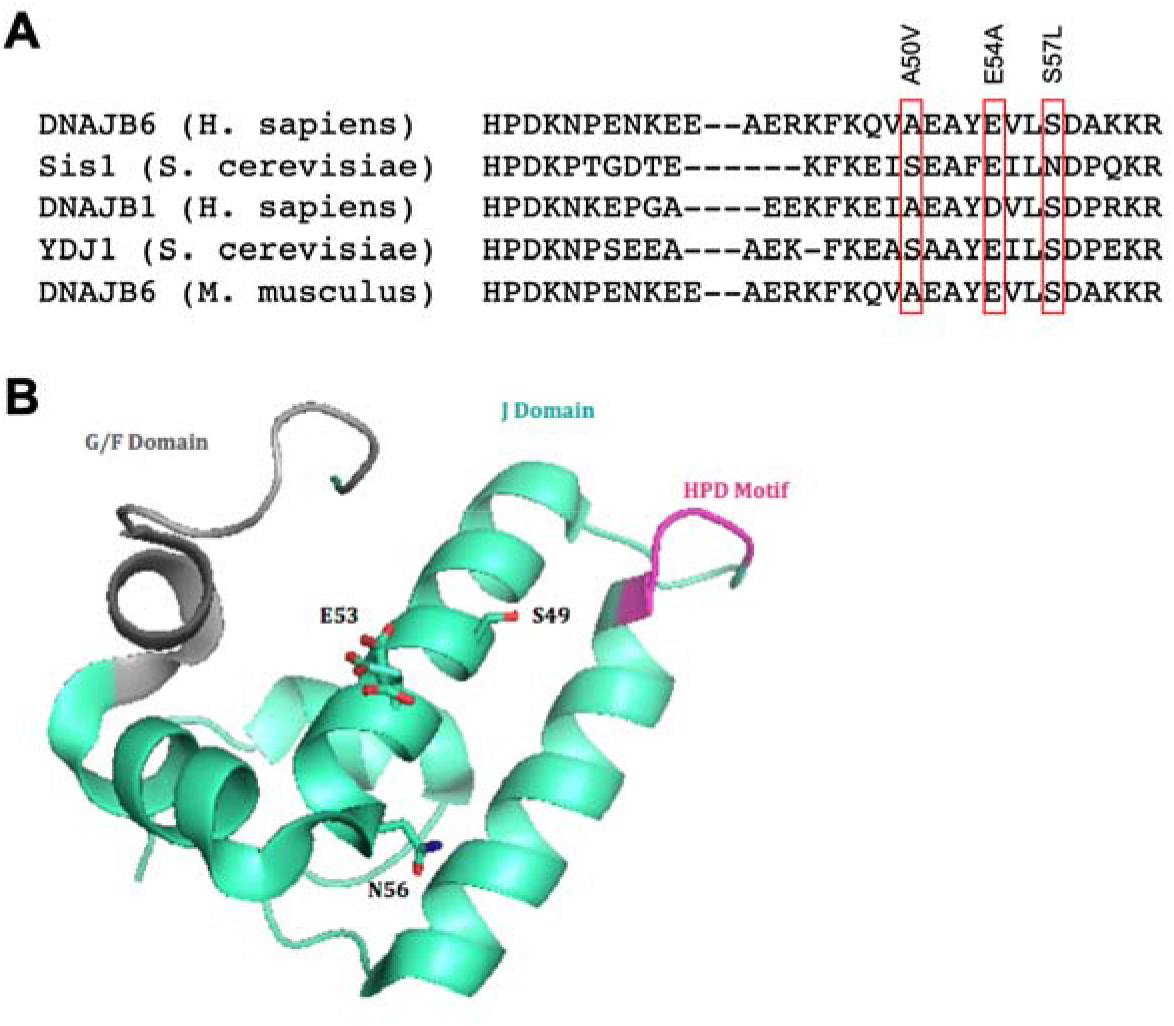
Novel LGMDD1 disease-associated mutations were identified in the J-domain of DNAJB6. (A) Amino acid sequence comparison of DNAJB6 and orthologs, including the yeast Hsp40, Sis1, aligned using ClustlW2. (B) Protein structure identifying homologous LGMDD1-associated mutations in the Sis1 J domain.

### Homologous mutations in the J domain of Sis1 differentially impair [PSI+] prion propagation

To assess the implications of these disease-causing variants on chaperone function, we turned to prion propagation models in yeast. Such models have been used extensively to study chaperones and understand the deleterious effects of disease-associated mutations on protein folding (40–46). In yeast, prions are naturally occurring, self-propagating protein structures that require chaperone networks for their efficient propagation (14). Defects in chaperone function disrupt this process and, therefore, prion propagation (27,33–36,47,48). In order to ask whether these disease variants disrupt client processing, we studied two known Sis1 substrates: the translation termination factor Sup35, which can aggregate to form the [*PSI*^+^] prion, and Rnq1, which can aggregate to form the [*RNQ*^+^] prion (14,29).

In an effort to understand the effect of homologous J domain mutations on [*PSI*^+^] prion propagation we performed experiments utilizing strains harboring the *ade1-14* allele that has a premature stop codon in the *ADE1* open reading frame (14). When Sup35 misfolds and aggregates in the [*PSI*^+^] state, nonsense suppression of the premature stop codon in *ade1-14* occurs and the adenine biosynthesis pathway is completed. Thus, [*PSI*^+^] colonies are white in color and can grow on media lacking adenine. Conversely, if cells are [*psi*−], adenine biosynthesis is incomplete, resulting in cells that are unable to grow in media lacking adenine and an accumulation of red pigment in cells grown on rich media. This colorimetric phenotypic assay has been utilized to demonstrate the effect of LGMDD1-causing G/F domain mutations on [*PSI*^+^] propagation (37). Here, we make use of four different [*PSI*^+^] strains (weak, strong, Sc37, and Sc4) (49,50). Each prion strain is a different self-propagating conformation of the same protein sequence and they are differentiated by the strength of the nonsense suppression phenotype and stability of their mitotic inheritance (49,51–53). Yeast containing *ade1-14* that propagate stronger [*PSI*^+^] strains (strong and Sc4) display higher nonsense suppression and are lighter in color. Yeast propagating weaker [*PSI*^+^] strains (weak and Sc37) are darker in color due to decreased nonsense suppression (better translation termination). Sis1 is required for yeast viability (54). Therefore, to perform these experiments, we used *sis1*Δ yeast (covered by a plasmid expressing Sis1) propagating different [*PSI*^+^] strains (weak, strong, Sc37, and Sc4) and replaced *SIS1* with homologous LGMDD1 mutant *SIS1* constructs or wild-type *SIS1* (as a control) (Fig 2A). In doing so, we observed viability was not affected in these strains expressing the mutants.

**Fig 2.**
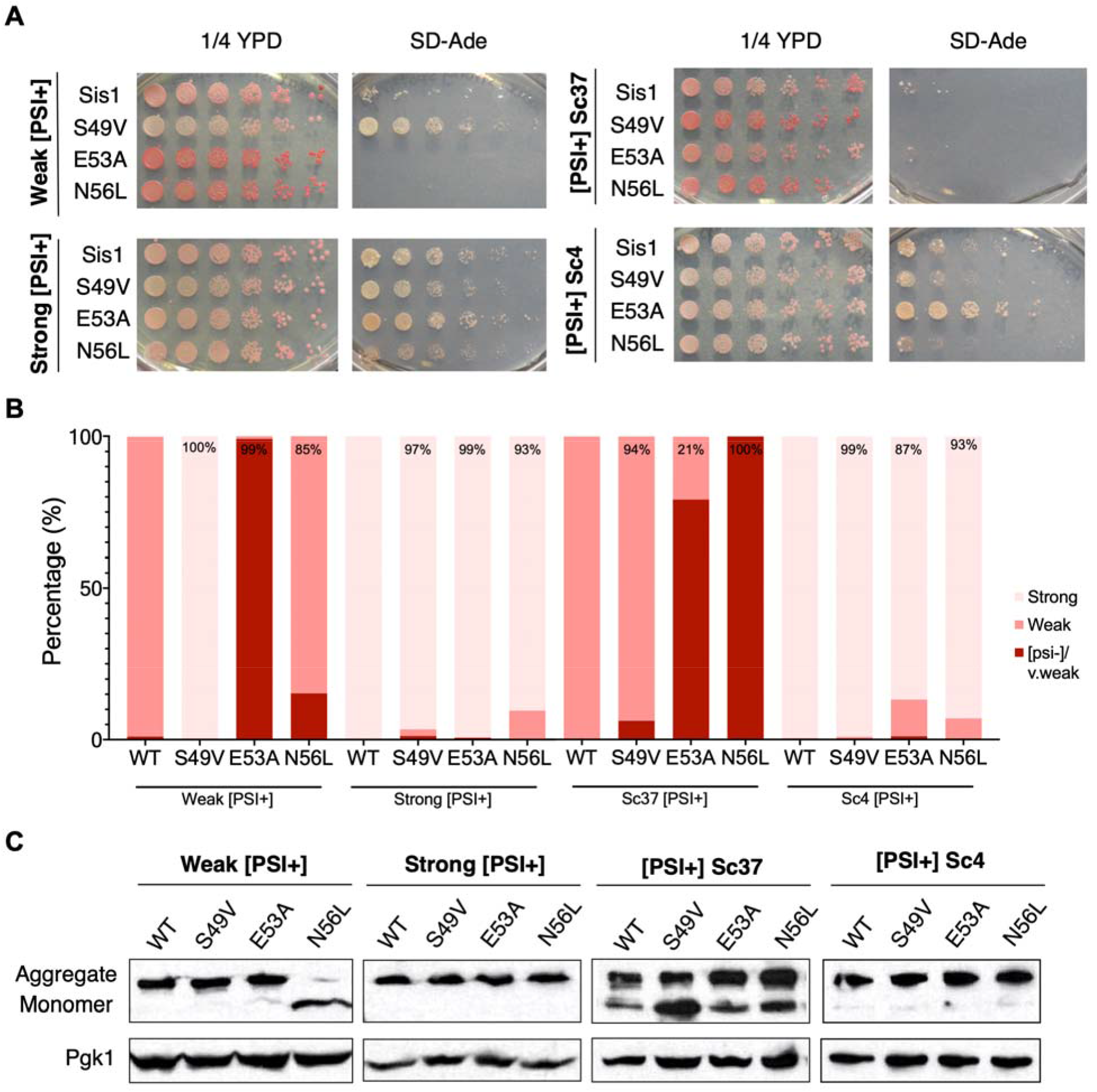
Homologous LGMDD1-associated mutations in the Hsp40 Sis1 differentially impair propagation of [PSI^+^] strains. (A) Yeast 74-D694 *sis1*Δ [*PSI*^+^] strains expressing wild-type *SIS1* or mutated *SIS1* constructs were serially diluted 5-fold and spotted onto 1/4 YPD media and SD-Ade to monitor nonsense suppression of the *ade1-14* allele (n=3). (B) [*PSI*^+^] colonies expressing the indicated constructs were isolated, grown in liquid YPD overnight, and plated on large 1/4 YPD spread-plates. An average of 626 colonies were counted and scored for strong [*PSI*+], weak [*PSI*+] or [*psi*−]/very weak phenotype. Data were collected from three separate biological replicates. (C) Representative western blot of yeast *sis1*Δ [*PSI*+] strains expressing wild-type *SIS1* or mutated *sis1* constructs. Cells were lysed and subjected to a boiled gel assay in order to display the amount of aggregated and monomeric Sup35 with antibodies against Sup35. Pgk1 is shown as a loading control. While we observed some reproducible differences in the percentage of monomeric Sup35 as compared to wild-type, we found these differences were not statistically significant. Standard protein markers are inappropriate for this particular assay and therefore are not shown (n=4-7). All samples were analyzed under the same experimental conditions.

Interestingly, we found that expression of Sis1-S49V in the weak [*PSI*^+^] strain, but not the Sc37 [*PSI*^+^] strain, altered the weak strain phenotype, as indicated by the lighter color on rich media and robust growth in selective media as compared to wild-type (Fig 2A). In comparison to the other J domain mutants, the stronger nonsense suppression phenotype we observed is unique to expression of Sis1-S49V in the weak [*PSI*^+^] strain. This suggests altered processing of Sup35 by Sis1-S49V occurs with conformational specificity. We questioned whether expression of Sis1-S49V could have altered the conformation of Sup35 in weak [*PSI*^+^] in a manner that leads to the propagation of a strong [*PSI*^+^] strain. Intrigued by this result, we introduced pRS316-Sis1 (wildtype) to this strain and re-assessed its [*PSI*^+^] phenotype (S1 Fig). We found this reverted the phenotype and was comparable to wild-type controls, suggesting that Sis1-S49V expression does not alter the [*PSI*^+^] strain. Previous work demonstrated that a shortage of cytosolic Sis1 may lead to more efficient transmission of propagons to daughter cells thus “strengthening” the prion phenotype (55). Thus, we hypothesize this may occur when Sis1-S49V is expressed in a weak [*PSI*^+^] strain.

This assay also showed that the expression of either Sis1-E53A or Sis1-N56L altered the propagation of the weaker [*PSI*^+^] strains (weak and Sc37), albeit in a different manner and to varying degrees. This is indicated by the phenotypic change towards a red color (less nonsense suppression) on rich media and lack of growth on media lacking adenine (Fig 2A). Similarly, when we introduced a wild type copy of *SIS1* to these phenotypically red strains, we identified a partial rescue of the weak [*PSI*^+^] phenotype (S1 Fig) indicating these Sis1 mutants, although phenotypically [*psi*−], result in propagation of cryptic [*PSI*^+^] propagons (56).

Complementarily, we used this assay quantitatively by phenotypically scoring large numbers of colonies on rich media spread plates (Fig 2B). This assay allows for a more detailed understanding of any changes in [*PSI*^+^] propagation. We scored colony color as light pink (strong [*PSI*^+^]), dark pink (weak [*PSI*^+^]) or red (very weak [*PSI*^+^] or [*psi*−]) in yeast expressing wild type Sis1 or the homologous LGMDD1 mutants. Indeed, through this assay we found that the expression of these mutants phenotypically altered prion propagation and identified a phenotypic variation that was not previously appreciated in the spotting assay. We observed a wider range of phenotypic variation in the weaker strains than in the stronger [*PSI*^+^] strains, which rarely convert to a weaker phenotype (57,58). However, an increase in the amount of darker pink colonies was observed when Sis1-S49V and Sis1-N56L were expressed in the stronger [*PSI*^+^] strains. Similar results were observed when Sis1-E53A and Sis1-N56L were expressed in the Sc4 [*PSI^+^*] strain (Fig 2B).

To further explore these results with respect to the solubility of the prion protein, we performed boiled gel assays to examine the relative levels of aggregated and monomeric Sup35. This assay consists of loading unboiled samples onto a sodium dodecyl sulfate– polyacrylamide gel electrophoresis (SDS-PAGE) gel. By doing so, non-denatured aggregates are too large to enter the resolving gel while monomeric Sup35 readily enters the gel due to its size and lack of aggregation. Halfway through electrophoresis, the gel itself is boiled and then run again, which allows previously aggregated protein to enter the resolving gel. The boiled gel assay provides a clear separation of monomeric and aggregated species of Sup35 and is easily detected through subsequent western blots (59). In order to examine the distribution of Sup35 between the aggregated and monomeric forms, we grew cultures from colonies representative of the phenotypic distribution observed in our spread plate assay.

Expressing homologous J domain mutations in the strong or Sc4 [*PSI*^+^] strains did not alter Sup35 distribution nor phenotype significantly (Fig 2C). Interestingly, the distribution of Sup35 between monomer and aggregate does not always correlate to phenotype in the weaker [*PSI*+] strains (many replicates of these assays were performed to verify reproducibility). Of note, we previously described a chaperone mutant that demonstrated differences in the ability to relate SDS-resistant or small protein aggregates to phenotype (56). Moreover, we found that expressing Sis1-S49V in the weak [*PSI*^+^] strain led to Sup35 being found mostly in the aggregated state (Fig 2C), coinciding with our phenotypic assay. With regards to expression of Sis1-E53A in the weak [*PSI*^+^] strain, we were surprised to find little Sup35 in the monomeric fraction, while the expression of Sis1-N56L results in most of the Sup35 in the monomeric fraction. Interestingly, expressing Sis1-S49V, Sis1-E53A or Sis1-N56L in the Sc37 [*PSI*^+^] strain showed a reproducible yet not statistically significant increase in monomeric Sup35 while maintaining a modest level of aggregated Sup35. Thus, there are differences with respect to the interaction between the Sis1 mutants and Sup35 in the weaker [*PSI*+] strains and how propagons and large aggregates correlate to phenotype.

### Homologous mutations in the J domain of Sis1 differentially impair [RNQ+] prion propagation

We then asked whether mutations in the Sis1 J domain would also affect another known client of Sis1, the [*RNQ*^+^] prion. In a manner similar to that of the previous experiments, we utilized various [*RNQ*^+^] prion strains (*rnq*−, low, and very high) and expressed each of our Sis1 mutants. Here, we observed reduced growth when Sis1-S49V was expressed in [*psi*−] strains that we did not observe in the same yeast strains harboring the [*PSI*+] prion. Thus, we performed spottings (using 5-fold serial dilutions) and confirmed that expression of Sis1-S49V, but not Sis1-E53A or Sis1-N56L, resulted in a growth defect (Fig 3A). This effect was only observed when wild-type Sis1 was removed through plasmid shuffle and occurred when Sis1-S49V was expressed in either [*RNQ*^+^] or [*rnq*−] strains. This indicated the defect in growth was independent of [*RNQ*^+^] and it was possibly impacted by the loss of [*PSI*^+^]. Because Sis1 is essential for yeast growth, mutations in Sis1 have been shown to affect viability (20,54,60,61).

**Fig 3.**
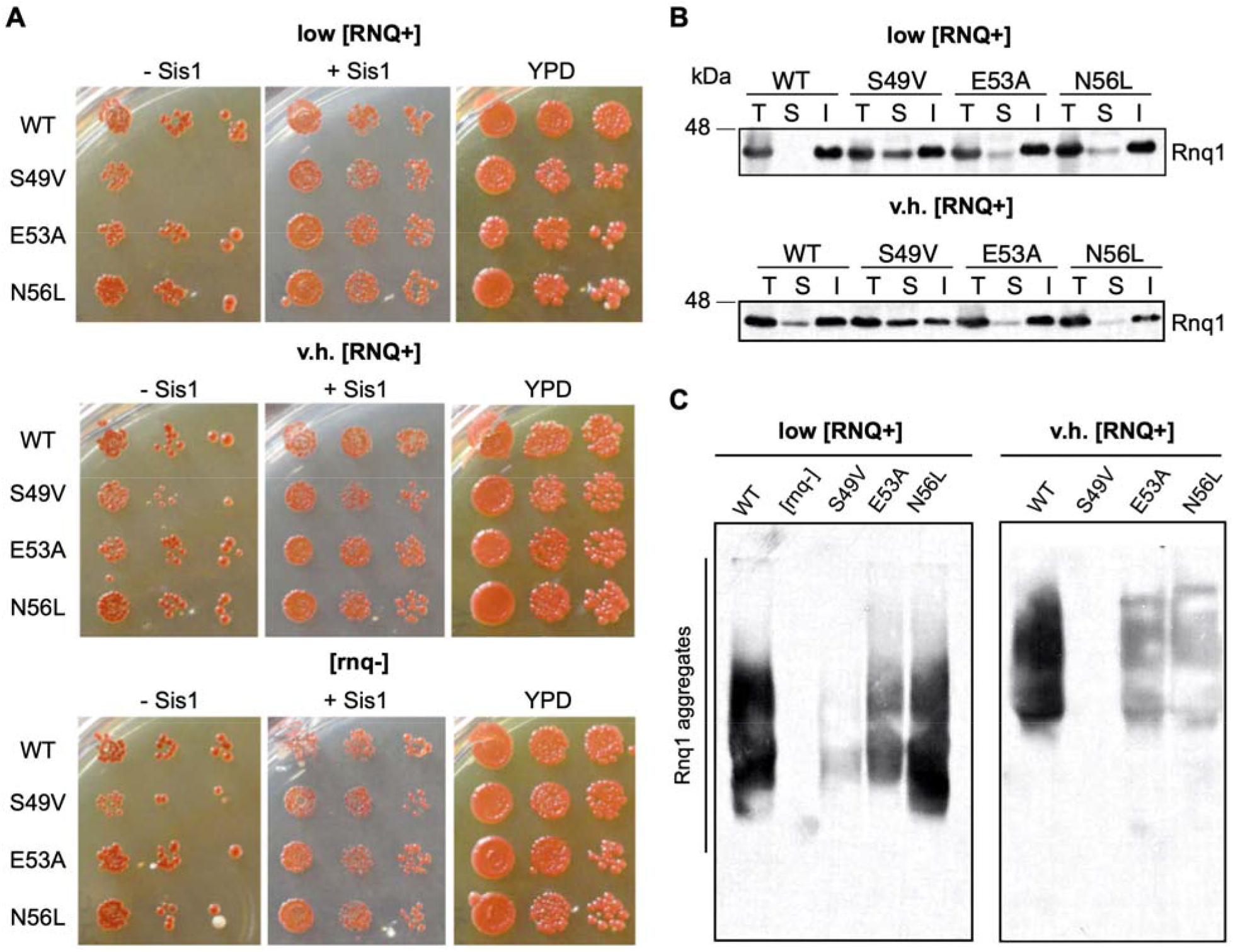
Homologous LGMDD1-associated mutations in the J domain of Sis1 differentially impair propagation of [RNQ^+^]. (A) Yeast *sis1*Δ cells harboring the indicated constructs and propagating low [*RNQ*^+^], very high [*RNQ*^+^] or [*rnq*−] were serially diluted 5-fold and spotted onto media in order to express the indicated *SIS1* constructs and select for either loss of wild-type Sis1 (−Sis1) or co-expression of wild-type Sis1 (+Sis1). Cells were also spotted on rich media (YPD) as a control for overall growth (n=3). (B) Sedimentation assay to separate Rnq1 into soluble and insoluble fractions with expression of the indicated constructs (n=6). Total (T), soluble (S) and insoluble (I) fractions were subjected to SDS-PAGE and western blot probing for Rnq1. (C) SDD-AGE assay for the indicated constructs in low [*RNQ*^+^] and very high [*RNQ*^+^] strains (n=4-6). All samples were analyzed under the same experimental conditions.

Unlike the nonsense suppression assay that discerns [*PSI*^+^] strains, strains of the [*RNQ*^+^] prion are characterized by the rate at which they induce the formation of [*PSI*^+^], ranging from low to very high rates (62–65). Therefore, we performed two different biochemical assays to assess [*RNQ*^+^] propagation. First, we performed a sedimentation assay (59) which consists of separating the soluble and insoluble protein fractions using ultracentrifugation to assay the relative solubility of Rnq1. These fractions are subjected to SDS-Page and probed with an antibody specific to Rnq1. We performed this assay using two [*RNQ*^+^] strains, low [*RNQ*^+^] and very high [*RNQ*^+^]. Using this assay, we identified a defect in [*RNQ*^+^] propagation when Sis1-S49V is expressed in both low [*RNQ*^+^] and very high [*RNQ*^+^] strains (Fig 3B). In comparison, there is only a slight change in Rnq1 solubility when Sis1-E53A or Sis1-N56L are expressed, indicating that J domain variants differentially impair processing of client conformers, albeit not completely.

Furthermore, while the sedimentation assay allows us to detect pelletable aggregates, these aggregates differ in their SDS sensitivity. In an effort to elucidate whether Rnq1 aggregate size or SDS resistance was altered by expression of the mutants, we performed a semi-denaturing detergent agarose gel electrophoresis (SDD-AGE) (Fig 3C). This assay allows for the visualization of SDS-resistant aggregates such as aggregated prion conformers (66). Strikingly, we observed a dramatic loss of SDS-resistant Rnq1 aggregates in the low [*RNQ*^+^] and very high [*RNQ*^+^] strains when Sis1-S49V was expressed. We also observed a slight reduction in SDS-resistant Rnq1 aggregates in low [*RNQ*^+^] strains expressing Sis1-E53A, while low [*RNQ*^+^] strains expressing Sis1-N56L appeared unaffected. In addition, very high [*RNQ*^+^] strains expressing Sis1-E53A or Sis1-N56L showed a decreased in SDS-resistant Rnq1 aggregates. Altogether, these biochemical assays present varying degrees of altered [*RNQ*^+^] propagation that occur in a conformer-specific manner when disease-associated J domain mutants are expressed in these strains.

### Disruption in prion propagation is due to a defect in substrate refolding

We observed different degrees of alteration in prion propagation due to the expression of the homologous LGMDD1 mutations in the J domain, which vary depending on the client and its conformation. To further understand the effect of these disease-associated mutations, we assessed how they impact protein folding of a non-prion substrate *in vivo*. Thus, we utilized the well-established firefly luciferase (FFL) refolding assay (67,68) (Fig 4). FFL is a non-prion substrate of the Hsp40/Hsp70/Hsp104 chaperone machinery and is denatured in cells by heat shock. When FFL misfolds and aggregates it requires the Hsp40/Hsp70/Hsp104 chaperone machinery to efficiently refold. Since Hsp104 is required for efficient refolding of FFL, we used a *hsp104*Δ strain expressing FFL as a negative control. We observed a significant defect in FFL refolding when Sis1-S49V was expressed in either a [*PSI*^+^] [*RNQ*+] (Fig 4) or [*psi*−] [*rnq*−] strain (S2A Fig), but not when other J domain variants are expressed. Of note, we did not observe a decrease in Sis1-S49V steady-state levels when this construct was expressed in Sc37 [*PSI*^+^] strains (S3 Fig). We subjected other known G/F domain mutants expressed in a chimeric protein where the G/F domain of Sis1 is replaced by that of DNAJB6, referred to as SDSS (37), to this assay and saw no significant differences in luciferase refolding as compared to wild type (S2B Fig). Hence, these results highlight the importance of substrate recognition – both with the client itself, as well as conformer specificity. Most importantly, these findings emphasize the distinction of assessing native chaperone clients as these mutants displayed varying degrees of altered prion propagation in our assays. Given this specificity, care should be taken when using certain assays, or a limited number of them, when trying to understand and confirm whether a potentially pathogenic chaperone variant is disease-causing or not.

**Fig 4.**
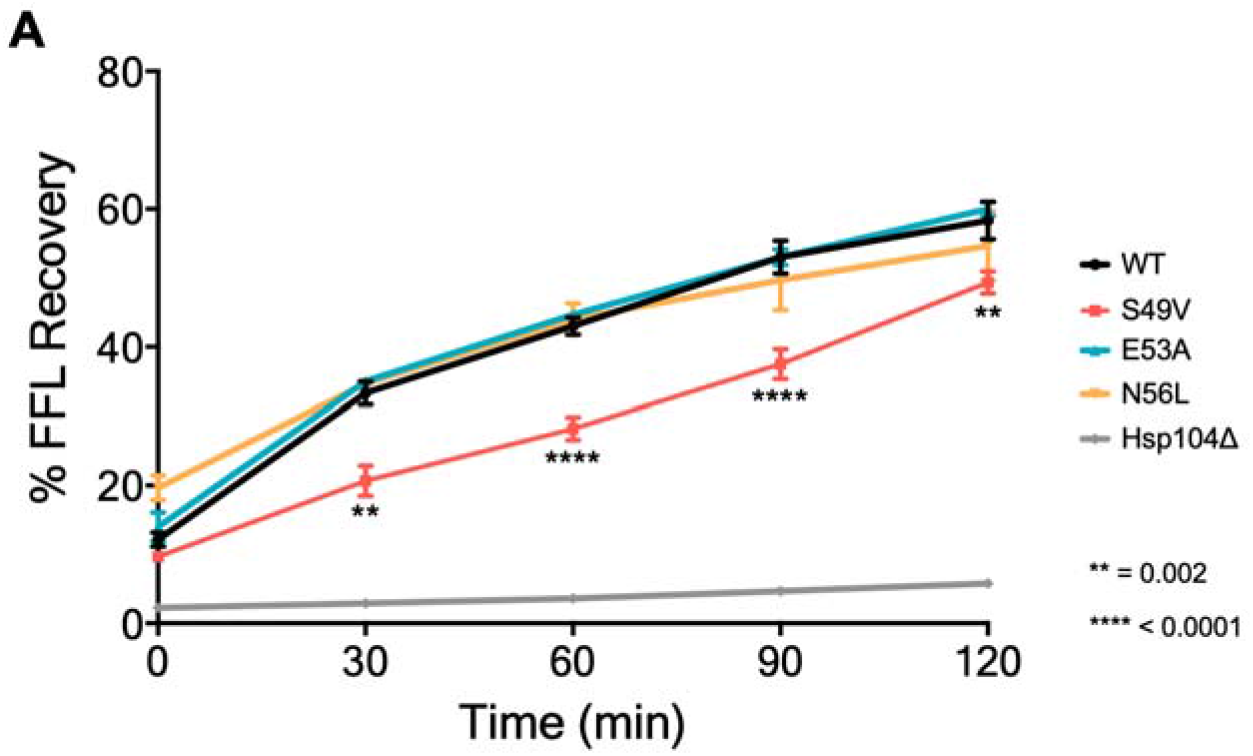
A homologous LGMDD1-associated mutant in Sis1 exhibits impaired substrate refolding. (A) The capability for refolding luciferase was measured in Sc37 [*PSI*^+^] *sis1*Δ yeast strains harboring the indicated construct along with a plasmid expressing firefly luciferase (FFL). Yeast were normalized, treated with cycloheximide and subjected to heat shock at 42°C for 22 minutes, followed by recovery at 30°C. Luminescence was measured at the indicated timepoints during recovery and normalized to luminescence of samples without heat shock treatment. The results represent the amount of luciferase refolding plotted as the percentage of recovery and represented as mean SEM (n=6). Each triplicate was averaged, (two-way ANOVA was performed; **p indicates a significant difference in Sis1-S49V FFL refolding relative to WT (p= 0.002) at the indicated timepoint; ***p indicates a significant difference in Sis1-S49V FFL refolding relative to WT at the indicated timepoint (p<0.0001)). All samples were analyzed under the same experimental conditions.

### Stability of a mutated Hsp40 chaperone is rescued by expression of the [PSI+] prion

While assessing the effect of the J domain LGMDD1 mutations on [*RNQ*^+^] propagation we identified a growth defect that only occurred with expression of the Sis1-S49V variant (Fig 3A). Interestingly, we observed this defect when Sis1-S49V was expressed in [*RNQ*^+^] [*psi*−] strains, but not when expressed in [*PSI*^+^] strains (Figs 2A and 3A). Because *SIS1* is essential (20,54,60,61), we were curious whether this growth defect was due to any change in steady-state expression levels. We assessed steady state Sis1 levels by western blot (Fig 5A) and found a significant decrease in the level of Sis1-S49V in various [*rnq*−] and [*RNQ*^+^] strains (Fig 5B). We hypothesize that this decrease in steady-state levels is responsible for the growth defect.

**Fig 5.**
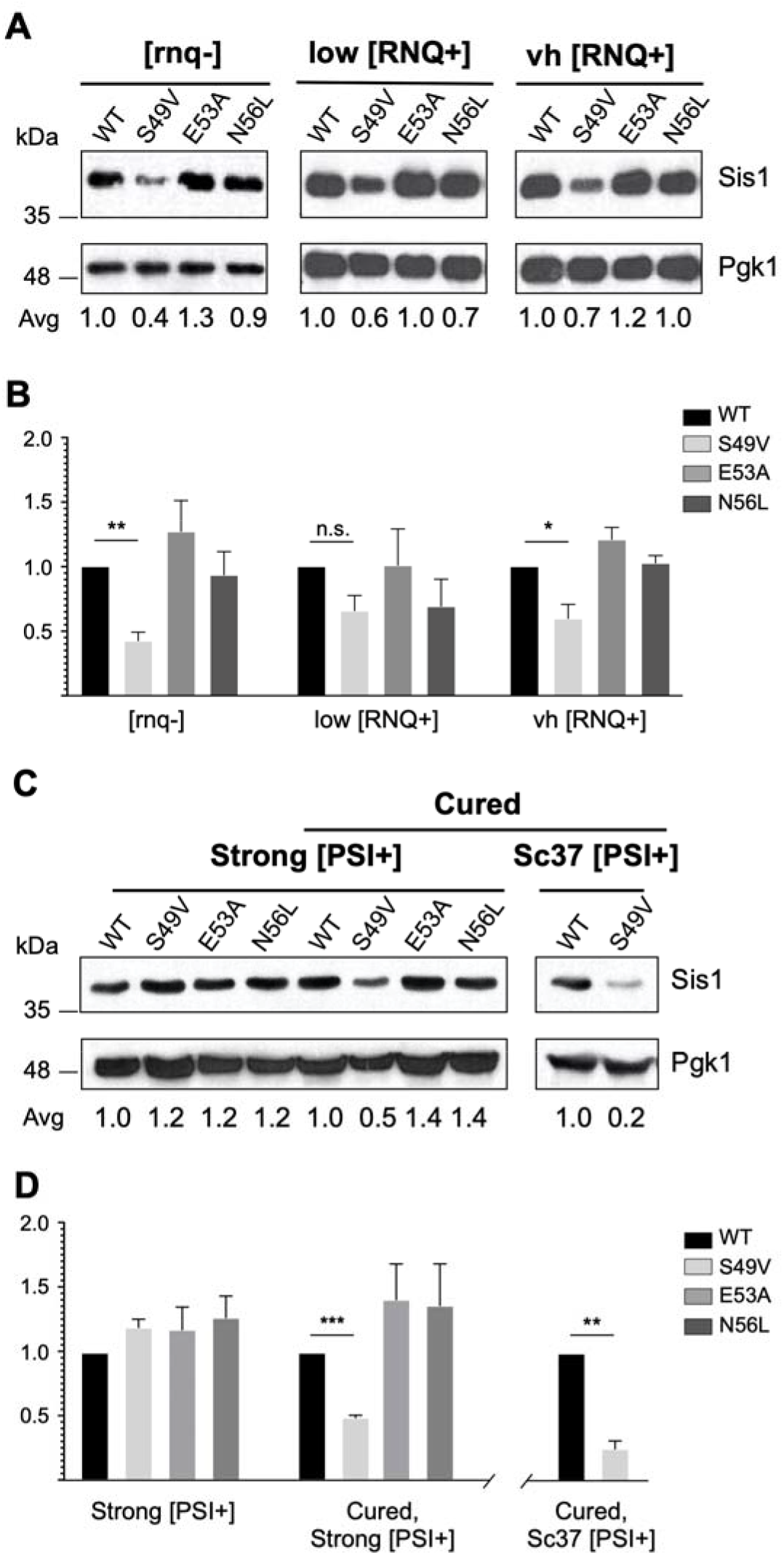
The LGMDD1-associated J-domain mutation Sis1-S49V has reduced steady state levels in the absence of [PSI^+^]. (A) Representative western blot showing expression of the indicated constructs in [*rnq*−], low [*RNQ*^+^], and very high [*RNQ*^+^] strains. (B) Results represent the quantified protein levels of Fig 5A displayed as their ratio relative to WT. (C) Representative western blot of strong [*PSI*^+^] *sis1*Δ yeast expressing the indicated construct (left). The specified strong [*PSI^+^*] and Sc37 [*PSI^+^*] strains expressing the indicated constructs were passaged twice on 3mM guanidine hydrochloride (GdnHCl) (cured), then grown on complete media without GdnHCl and lysed for SDS-PAGE followed by western blot (right). Pgk1 served as a loading control. (D) Results represent the quantified protein levels of Fig 5C displayed as their ratio relative to WT. For all Western Blot analyses, intensities were quantified and plotted as the mean SEM (n=3). Independent experiments indicating a significant difference in Sis1 expression levels relative to WT (paired t-test; n.s. nonsignificant; *p<0.05; **p<0.01; ***p<0.001). All samples were analyzed under the same experimental conditions.

To confirm this decrease is due to a lack of [*PSI*^+^] presence, we set out to recapitulate this phenomenon by curing [*PSI*^+^] strains of its prion form. Curing of [*PSI*^+^] was performed by growth on media containing guanidine hydrochloride (GdnHCl), which inactivates the ATPase activity of Hsp104 and hinders the replication of [*PSI*^+^] seeds (69). After curing both a strong [*PSI*^+^] and a Sc37 [*PSI*^+^] strain (each expressing Sis1-S49V) we performed western blots (Fig 5C) and identified a significant decrease in Sis1 steady-state levels (Fig 5D). This decrease was not observed with expression of wild type Sis1 nor the other LGMDD1 J domain variants. The decrease in steady-state levels recapitulates what we observed in [*psi*−] strains expressing Sis1-S49V (Figs 5A and 5B). This striking result suggests the Sis1 client, Sup35, in its aggregated and [*PSI*^+^]-propagating form, interacts with this particular Hsp40 mutant in a way that stabilizes the steady-state level of Sis1-S49V. Of note, this decrease in Sis1 steady-state levels has not been observed in the context of LGMDD1-associated mutations found in the G/F domain. This interesting finding is the first known instance wherein a potentially unstable mutant chaperone is stabilized by the presence of a specific client protein in its aggregated form.

## Discussion

LGMDD1 is a myopathy historically characterized by mutations within the G/F domain of the chaperone DNAJB6 (6–9,11). Here we used the yeast model system, a homologous chaperone, and native chaperone clients to demonstrate that recently reported mutations in the J domain of DNAJB6 alter canonical chaperone function. Specifically, expression of homologous disease-associated mutations in the yeast Hsp40, Sis1, identified defects in prion propagation to varying degrees depending on the disease-associated variant, the client protein and its conformation, as well as a significant defect in refolding of a non-prion client protein. We conclude these mutations in the J domain of Sis1 lead to aberrant chaperone function, altered protein homeostasis, and potentially drive a variety of defects in chaperone machinery which may contribute to pathogenesis.

The function and interaction of the various Hsp40 domains has been a topic of interest for many years (20,21,70–75). Surprisingly, colleagues identified LGMDD1 patients with novel variants in the J domain of DNAJB6, a disease previously characterized by mutations in the G/F domain (12,13). As previously mentioned, LGMDD1 disease-associated mutations in the G/F domain have been found to impair the processing of specific client conformers. These findings highlighted the G/F domain as having an important role in substrate regulation and conformer selectivity (37). By contrast, the J domain of Sis1 has a known role in regulating Hsp70 ATPase activity (76).

Here we demonstrate a novel finding by which disease-associated missense mutations in the J domain of an Hsp40 result in the chaperone having client and conformer specificity. Strikingly, this fits with previous structural studies which have highlighted the importance of the intramolecular interaction between the J and G/F domains (21). Specifically, a particular amino acid, E50, in the Sis1 J domain has been shown to interact with the EEVD(HSP70) motif, an interaction required for both chaperones to partner in protein refolding (21). Moreover, this key amino acid (E50) forms a salt bridge with R73 which is found in the Sis1 G/F domain. We and others (12,21) believe this interaction is crucial and of functional importance for the stability and interaction between both domains and, ultimately, the interaction between Hsp40 and Hsp70.

In the context of the LGMDD1-associated mutants found in the J domain, all variants appear to affect prion propagation to varying degrees depending on client and conformation specificity. Notably, the homologous Sis1-S49V variant appears to have the most deleterious effect on the chaperone machinery and its function, probably due to its proximity to the E50:R73 salt bridge. This would also explain why only Sis1-S49V presented with defective substrate refolding in our luciferase refolding assay. Overall, we hypothesize that the steric hindrance caused by these missense mutations may obstruct the intramolecular interaction or communication between the J and G/F domains. Moreover, these findings support the idea that it may be the interaction between these two domains that is crucial for the specificity and processing of Hsp40 client proteins.

By assessing the propagation of various Sis1 clients, we found that there are differences in prion propagation, but that propagation is not completely abolished by expression of these mutants. When compared to LGMDD1-associated mutations in the G/F domain, for example, we observe variants in the G/F domain to have a stronger effect on prion propagation in certain assays, oftentimes demonstrating complete loss of prion propagation (37). As such, although these disease-associated mutations in the J domain disrupt prion processing and protein refolding, they do so in a manner that is slightly different from that of G/F domain mutants.

Given the limited knowledge of the clients of DNAJB6 in skeletal muscle and its role in disease, we hypothesize, as we did for the G/F domain mutations, that distinct variants may be associated with different levels of disease severity due to impairment of Hsp40 function to varying degrees, or in different ways. Nonetheless, we are cautious to imply that a more drastic loss-of-propagation effect would correlate with or be causative of a more severe disease pathogenesis, as we have observed different degrees of aberrant chaperone function in the context of these disease-associated variants that are greatly dependent on client and conformation specificity. It is not uncommon for LGMDD1-associated mutations to have an effect in some but not all assays or when assaying specific client proteins (12,37). Moreover, as we continue to understand and classify disease-associated variants as pathogenic, we should consider how variable clinical outcomes and disease severity may be influenced by factors such as variable expressivity and sex influences (11). Given these considerations, there is a need for simple models, such as the yeast system, by which we can understand if and how identified genetic variants affect chaperone function in order to better understand variable clinical outcomes within LGMDD1. Currently, we can only show that there are differences in adequate client processing and refolding although, undoubtedly, further *in vitro* studies are required to further elucidate the mechanistic cause of these functional defects and how they translate to pathogenesis.

Lastly, we identified a novel phenomenon by which a mutated Hsp40, Sis1-S49V, appears to have a decreased steady state level of protein expression. This is, to our knowledge, the first known instance where steady-state expression of a chaperone is dependent on stabilization by a known substrate. Although serendipitous, this finding not only highlights an important function of chaperone-substrate interaction, but also hints at an additional role that specific substrates might play in chaperone function and protein homeostasis overall. Indeed, there is still much to be learned about co-chaperones, their function, interactions between themselves and with clients, and their role in disease pathogenesis.

## Acknowledgements

We thank S. Liebman, J. Weissman and E. Craig for reagents. We also thank J. Daw for assistance with site-directed mutagenesis. We are appreciative of L. Dublin and A. Bhadra for helpful discussions and comments on the manuscript.

## Supporting information

**S1 Table.**
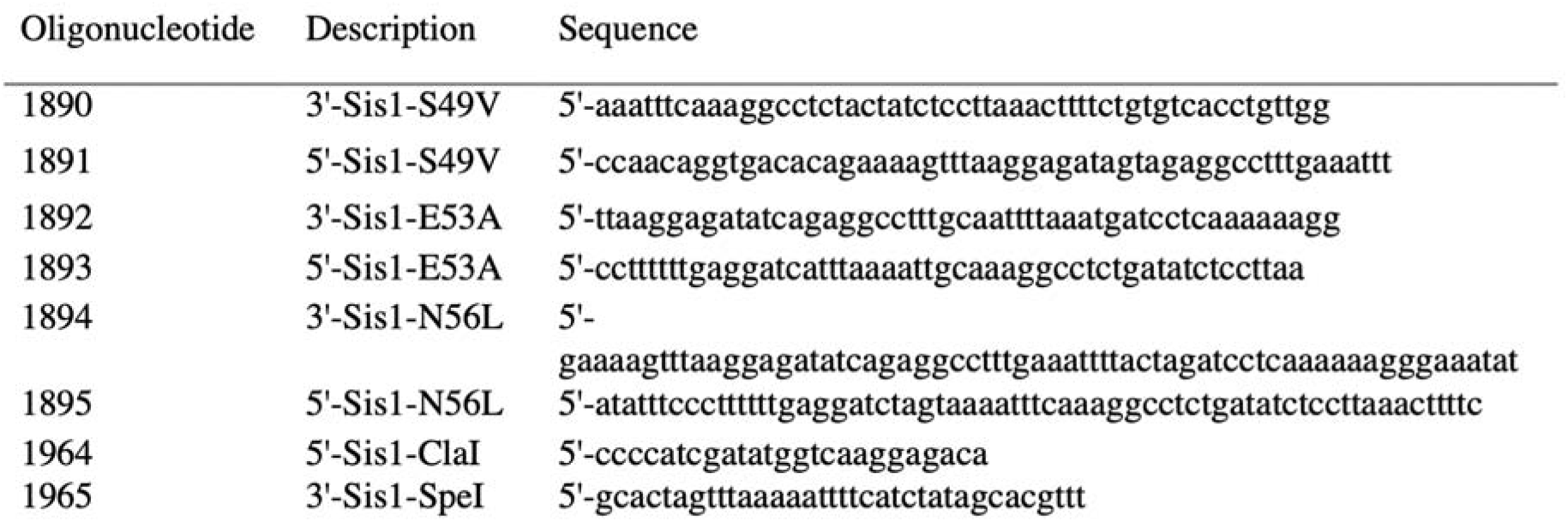
Oligonucleotides used in this study.

**S1 Fig.**
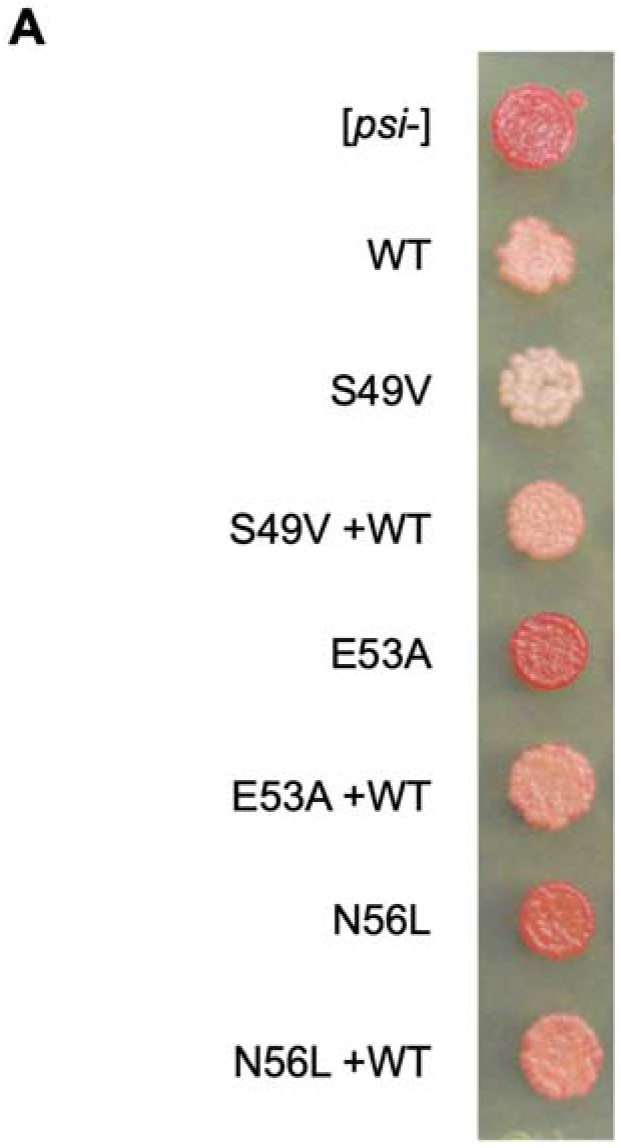
Novel LGMDD1 J-domain mutations in Sis1 differentially impair propagation of [PSI^+^] strains and are rescued by expression of an additional WT-Sis1 copy. (A) *sis1*Δ [*PSI*^+^] strains expressing a single copy of wild-type or mutated *SIS1*, and strains expressing an additional copy of wild-type *SIS1* (+WT) were spotted onto YPD media (n=3). All spottings, minus the top (labeled [*psi*−]) are weak [*PSI*^+^] strains expressing the indicated constructs.

**S2 Fig.**
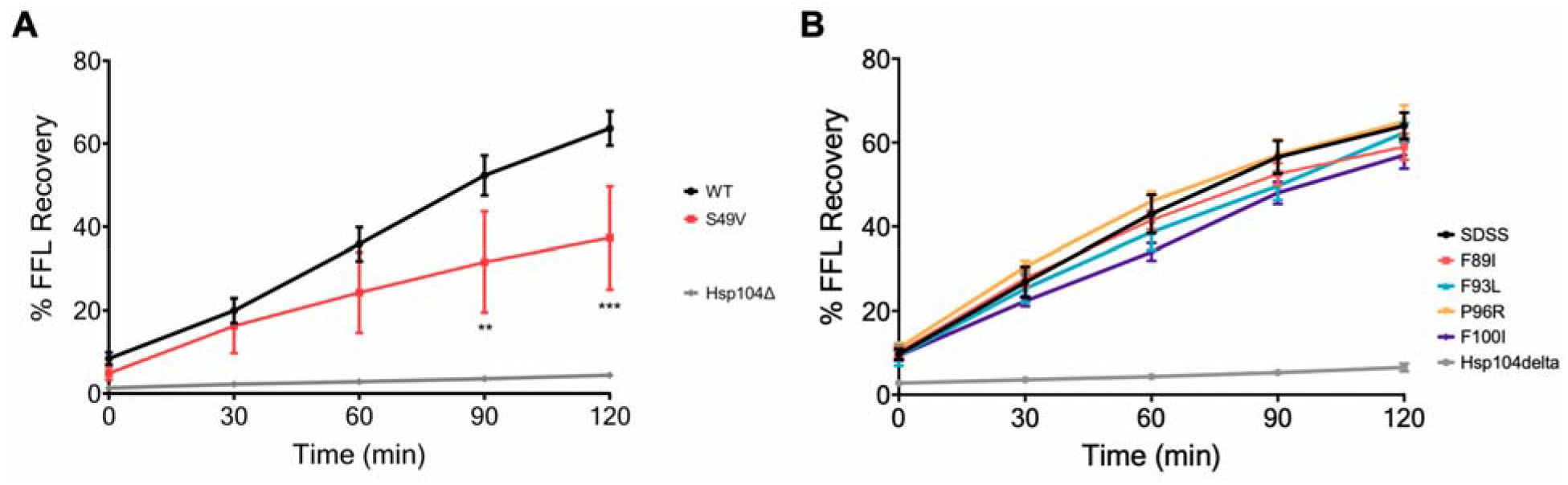
Sis1-S49V exhibits impaired substrate refolding while G/F domain mutants do not. (A) the capability for refolding luciferase was measured in [*rnq*−] *sis1*Δ yeast strains harboring the indicated construct along with a plasmid expressing luciferase. (B) same as in *A*, but in Sc37 [*PSI*^+^] strains expressing wild-type or the mutated chimeric construct *SDSS* constructs instead of Sis1. Yeast were normalized, treated with cycloheximide and subjected to heat shock at 42°C for 22 minutes, followed by recovery at 30°C. Luminescence was measured at the indicated timepoints during recovery and normalized to luminescence of samples without heat shock treatment. The amount of luciferase refolding is plotted as percentage of recovery and represented as mean SEM (n=6).

**S3 Fig.**
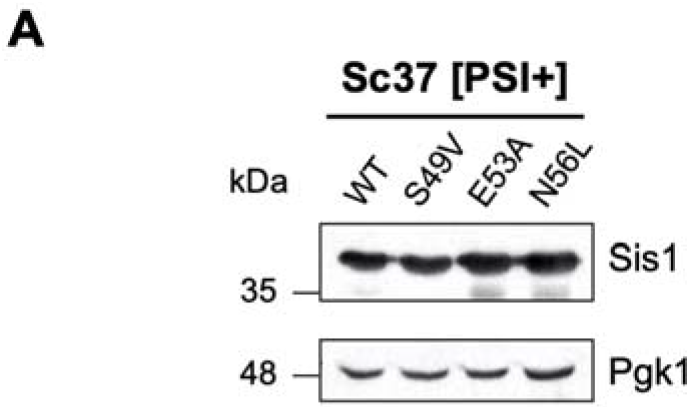
Sis1-S49V, does not show decreased steady-state levels when expressed in Sc37 [PSI^+^] strain. (A) Representative western blot showing expression of Sis1 in Sc37 [*PSI*^+^] *sis1*Δ yeast strains harboring the indicated construct (n=3). Pgk1 is shown as a loading control. All samples were analyzed under the same experimental conditions.

